# Diet-induced changes in the colonic microenvironment reduce polyp development in APC^Min/+^Msh2^-/-^ mice by triggering ER stress

**DOI:** 10.1101/2025.03.13.643056

**Authors:** María López Chiloeches, Antoaneta Belcheva

## Abstract

CRC development is a complex disease driven by the interplay between genetic mutations, diet, gut microbiota and inflammation. However, how these factors work together to exert their effect on disease initiation and progression is still unclear. In this study, we investigated the effects of a 10% low carbohydrate (10% LC) diet and a normal diet supplemented with 10% pectin from apple (ND+10% Pec) on CRC progression and colonic homeostasis. We showed that although these dietary regimens had differential effects on the gut microbiota and the production of butyrate, they both reduced CRC development in the APC^Min/+^Msh2^-/-^ mouse model. 10% LC diet strongly reduced the number of goblet cells while ND+10% Pec had no such effect, but both regimens induced upregulation of *Muc-2*. The changes in the gut microbiota, butyrate levels and increased Muc-2 expression, however, trigger endoplasmic reticulum (ER) stress. Our results show that a 10% LC diet triggered endoplasmic reticulum stress (ER stress) via induction of the PERK pathway, while ND+10% Pec also increased the splicing of the XBP1. In addition, both diets induced dramatic expression of the heat shock proteins Hsp27 and Hsp70, further resulting in increased barrier function. These events are likely to protect the cells and help them cope with environmental changes. However, the induced cellular stress conditions, known to suppress cell proliferation, could be a mechanism through which these regimens attenuate CRC.

## INTRODUCTION

Colorectal cancer (CRC) is a leading cause of cancer death. The mechanisms that drive CRC initiation and progression have been linked to the complex interplay between genetic mutations, the gut microbiota, diet and inflammation. The development of CRC is driven by stepwise accumulation of genetic alterations of oncogenes and tumour suppressor genes in colon epithelial cells (CECs) that affect the proliferation, differentiation, and the survival of these cells (Fearon 2011). The most common mutations associated with CRC initiation are in the adenomatous polyposis coli (APC) gene, as well as in genes implicated in the DNA mismatch repair (MMR) machinery (Fearon 2011). APC is a component of the Wnt/β-catenin signalling and plays a critical role in β-catenin degradation (Medema and Vermeulen 2011). MMR is the main pathway that repairs errors made during DNA replication, maintains genomic stability and signals apoptosis in response to DNA damage (Jiricny 2006, Poulogiannis, Frayling et al. 2010). The development of microsatellite instability and gene silencing due to promoter methylation are the two main mechanisms through which MMR-deficiency predisposes to CRC (Boland and Goel 2010). However, it has been shown that inactivation of MMR system in CECs causes increased proliferation that renders these cells highly susceptible to transformation by the gut microbiota and their metabolism of complex carbohydrates (Belcheva, Irrazabal et al. 2014). Therefore, the interaction between MMR genetic background and the colonic microenvironment is crucial in CRC initiation and development and more research is needed to elucidate this link.

The influence of microbial metabolism on CRC has been largely investigated (Frank, St Amand et al. 2007, Scanlan, Shanahan et al. 2008, Gao, Guo et al. 2015, Vipperla and O’Keefe 2016). Specifically, the gut microbiota carries out the fermentation of complex carbohydrates into short-chain fatty acids (SCFA). Butyrate is the principal SCFA and has diverse effects on intestinal epithelial cells. It is the main energy source and a potent signalling molecule that controls gene expression programs essential for the proliferation of the intestinal epithelium (Donohoe, Wali et al. 2012, Zeng, Lazarova et al. 2014). Butyrate also acts as an anti-inflammatory molecule. This role is mediated by the interaction of butyrate with G–protein-coupled receptor GPR109A, expressed on the surface of the intestinal epithelial cells and macrophages (Ganapathy, Thangaraju et al. 2013). Activation of this receptor in CECs leads to the production of IL-18 which is known to suppress inflammation (Singh, Gurav et al. 2014). In cancer cells, butyrate induces apoptosis via inhibition of the histone deacetylases (HDACs), which is a mechanism defining its cancer-protective effects (Donohoe, Holley et al. 2014, Encarnacao, Abrantes et al. 2015). However, butyrate may also act as an oncometabolite. It has been shown that in APC^Min/+^Msh2^-/-^ colon epithelial cells bacterially-produced butyrate stimulated the proliferation of CECs and fueled CRC development (Belcheva, Irrazabal et al. 2014). In this case, reducing the butyrate levels in the colon by treating the mice with antibiotics or a diet low in complex carbohydrates (7% LC diet) led to a significant reduction of polyps. The diverse effects of butyrate on CRC, therefore, may depend on the specific genetic background of the cells.

The colon epithelial cells are segregated from the cells of the gut microbiota by the inner and outer mucus layers. The major constituent of the inner mucus layer is Mucin 2 (Muc-2), which is secreted by a differentiated CEC subtype, the goblet cells (Cobo, Kissoon-Singh et al. 2015). The inner mucus layer is normally free of microorganisms, while the outer mucus layer is well-populated by members of the gut microbiota. The absence of complex carbohydrates and dietary fibre, which are the main energy source for the majority of the gut microbes, may lead to the expansion of mucus-associated microorganisms because these microbes can degrade Muc-2 to obtain energy (Desai, Seekatz et al. 2016). From another perspective, changes in the composition and availability of the macronutrients are associated with specific cellular responses such as endoplasmic reticulum (ER) stress, which allows the cells to adapt to the available metabolites (Wellen and Thompson 2010). Specifically, the unfolded protein response (UPR) plays a central role in the regulation of ER stress which is associated with nutrient adaptation. Three main proteins (PERK, IRE1 and ATF6) and their downstream signalling cascades help the cells to cope with the ER stress by inhibiting protein translation and regulating protein folding and degradation of misfolded proteins (Kaufman, Scheuner et al. 2002, Wellen and Thompson 2010). Changes in diet and associated alterations in the gut microbiota may also result in the induction of heat-shock proteins (HSPs), nicely reviewed by Arnal and Lallés (Arnal and Lalles 2016). HSPs are expressed by intestinal epithelial cells in response to microbial products, metabolites, oxidative stress, ER stress or/and inflammation (Liu, Musch et al. 2003, Rakoff-Nahoum, Paglino et al. 2004, Arvans, Vavricka et al. 2005, Tao, Drabik et al. 2006, Carlson, Vavricka et al. 2007). Nevertheless, the ability of diets to alter the composition and the metabolic activity of the gut microbiota remains an attractive means to attenuate and control disease. Indeed, most studies have focused on understanding the mechanisms through which specific diets and associated changes in the gut microbial composition affect the cells within the tumour. However, how the changes in the intestinal microenvironment affect intestinal homeostasis is unclear. Hence, a better understanding of the complex interplay between genetic mutations, diet and gut microbiota would shed light on our current knowledge of CRC-promoting mechanisms.

In this study, we investigated the effects of either a 10% LC or a normal diet supplemented with 10% pectin from apple (ND+10%Pec) on the composition of the gut microbiota, CRC progression and colonic homeostasis. We showed that although both dietary regimens reduced CRC development in the APC^Min/+^Msh2^-/-^ mouse model, they had differential effects on the gut microbiota and the production of butyrate, suggesting that these diets reduce CRC by different mechanisms. 10% LC diet strongly reduced the number of goblet cells while ND+10% Pec had no such effect, but both regimens induced upregulation of *Muc-2*. The changes in the gut microbiota, butyrate levels and increased Muc-2 expression, however, trigger ER stress. Interestingly, a 10% LC diet triggered endoplasmic reticulum stress (ER stress) via induction of the PERK pathway, while ND+10% Pec also increased the splicing of the XBP1. In addition, both diets induced overexpression of two heat shock proteins, Hsp27 and Hsp70, and increased barrier function.

## MATERIALS AND METHODS

### Mice and treatments

The Msh2^-/-^ mice have been previously described (Reitmair, Schmits et al. 1995). The APC^Min/+^ mice were purchased from Jackson Laboratory. All mice were on the C57BL/6J background and maintained under specific pathogen-free conditions at the animal facilities in the Biomedical Laboratory (University of Southern Denmark).

Msh2 heterozygous (Msh2^+/-^) mice are characterised by normal DNA repair and were used as controls in this study. APC^Min/+^Msh2^+/-^ and APC^Min/+^Msh2^-/-^ mice were generated and genotyped as described before (Belcheva, Irrazabal et al. 2014). The mice were fed with the 1320 Diet obtained from Altromin, Germany. This diet is designated as ND control. The 10% low-carbohydrate modification of this diet was also obtained from Altromin. The macronutrient composition of the used diets is shown in Supplementary Table 1. The 10% pectin diet was prepared by supplementating the ground ND with 10% pectin from apple (Sigma). This diet was prepared fresh every day. All dietary regimens were administered immediately after weaning for a period of 3 weeks. Some mice were also treated with metronidazole, provided in the drinking water at 1 g/L. The treatment started after weaning and lasted 3 weeks (Figure 1A).

**Figure 1.**
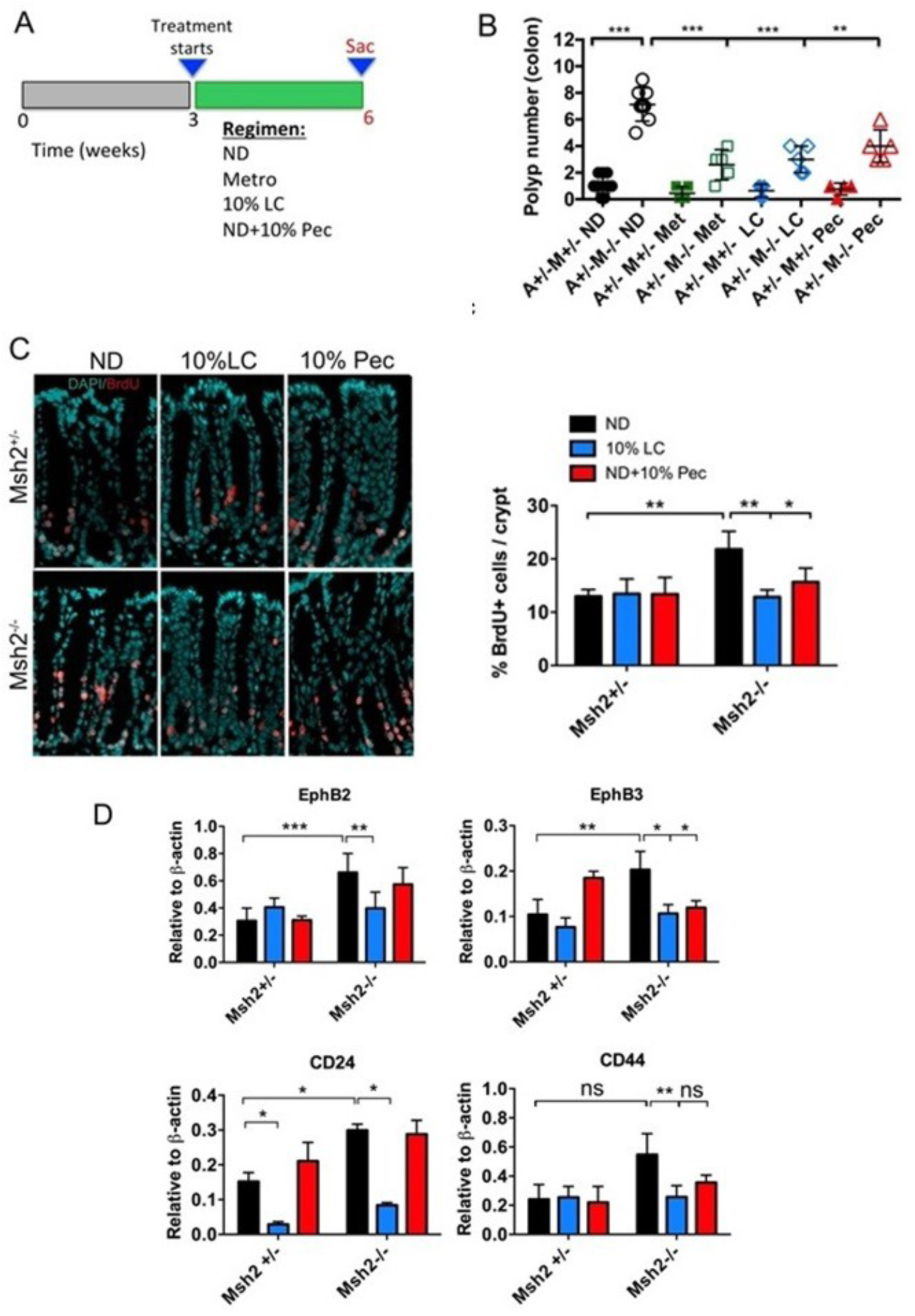
10% low carb (LC) and pectin-enriched (ND+10% Pec) diets reduce polyp formation and colonic cell proliferation in APC^Min/+^Msh2^-/-^ mice. (A) Schematic representation for administration of dietary regimens to APC^Min/+^Msh2^+/-^ and APC^Min/+^ Msh2^-/-^ mice. All treatments started 3 weeks after weaning and continued for 3 weeks. Mice were sacrificed at 6-week old and analysis were carried out. (B) Number of polyps counted in the colons of APC^Min/+^ Msh2^+/-^ (A+/-M+/-) and APC^Min/+^ Msh2^-/-^ (A+/-M-/-) mice under different treatments. Each symbol represents an individual mouse. (C) Assessment of the proliferation of colonic epithelial cells by immunostaining with bromodeoxyuridine (BrdU;red). Msh2^+/-^ and Msh2^-/-^ mice on indicated diets were injected with BrdU or PBS. After 2 hours the mice were sacrificed and BrdU positive cells were visualised by immunofluorescence. The magnification of the representative images is x40. Percentages of BrdU-positive cells per colonic crypt are represented in a histogram. (D) mRNA expression of stem cell markers EphB2, EphB3, CD24 and CD44 in colonic epithelial cells from Msh2^+/-^ and Msh2^-/-^ mice was quantified by qPCR. Data were normalized to β-actin. All data were analysed by unpaired *t-*test and *p*-values are also shown (* *p*-value ≤ 0.05, ** *p*-value ≤ 0.01, *** *p*-value ≤ 0.001).

### Analysis of colonic polyps

The APC^Min/+^Msh2^+/-^ and APC^Min/+^Msh2^-/-^ mice on different treatments were sacrificed and their colons were removed, cleaned with fresh PBS and opened longitudinally. The polyps present in the entire colon were identified and counted under a dissection microscope.

### Analysis of the gut microbiota

Stools were collected from the distal region of the colon and DNA was extracted with a QIAamp DNA Stool kit (Qiagen), according to the manufacturer’s instructions. DNA concentrations were measured in DeNovix DS-11 spectrophotometer and adjusted to a DNA stock with a final concentration of 20 ng/µl. Bacterial DNA (4 ng/µl) was analyzed by quantitative PCR (qPCR). The total bacterial abundance was quantified by universal 16S rRNA primers. The quantification of specific bacterial groups was carried out by previously described primers targeting these groups. The sequences of all primers used in this study can be seen in Supplementary Table 2. Fragments were amplified in StepOne Plus PCR System (Applied Biosystems) using the following program: 95 °C for 3 min, followed by 40 cycles of 95 °C for 30 seconds. The primer annealing optimal temperature was determined in this study (Supplementary Table 2) and was carried out for 30 seconds. Finally, the extension was at 72 °C for 30 seconds. Total bacterial abundance was calculated as the total number of *Eubacteria*, considering DNA concentration and sample weight. The relative abundance of each bacterial group was calculated as a percent of *Eubacteria* using the *ΔC_T_* method (*ΔΔC_T_*)(Yang, Chen et al. 2015).

### *In vivo* labelling of colon epithelial cells with BrdU

Mice on different dietary treatments were injected with 100 mg/kg BrdU sterile solution. BrdU was diluted in PBS at 10 mg/ml. After 2 hours, mice were euthanized, and dissected colons were prepared as Swiss rolls and frozen in OCT medium. 5 µm sections were cut and subsequently fixed in cold acetone for 7 min followed by hydration in PBS for 15 min. The tissue was then incubated in 1M HCl for 10 min on ice to break open the DNA structure of the cells. This was followed by incubation in 2M HCl for 10 min at room temperature. The slides were then moved to a 37°C incubator for 20 min. The tissue was neutralized by incubating the samples in 0.1M Borate buffer for 10 min at room temperature. The slides were washed 3 times 5 min each in PBS - 0.1% Triton X-100 and blocked in 1% BSA/10% goat normal serum /0.3M glycine in PBS-Tween 20 (0.1%) for 1 hour at room temperature. The slides were then processed by standard immunostaining procedure using BrdU antibody (Abcam, ab142567) at 1:100 for overnight at 4°C and secondary antibody Goat anti-rabbit IgG-Alexa 488 at 1:250 in PBS+ 1%BSA for 1 hour at room temperature. Finally, the nuclei were stained by 4’,6-diamino-2-phenylindole (DAPI) from Sigma, diluted at 1:1000 in PBS. The fluorescent imaging was performed with a Zeiss LSM510 confocal microscope at the Danish Molecular Biomedical Imaging Center (DaMBIC, University of Southern Denmark).

### Isolation of colon epithelial cells

Colon tissues from Msh2^+/-^ and Msh2^-/-^ mice were collected and pure intestinal crypts were isolated as follows. The colons were removed, flushed with PBS and cut into 0.5 cm pieces that were subsequently transferred in Ca^2+^/Mg^2+^ free PBS supplemented with 5 mM EDTA. The tissue was incubated for 20 min at 37 °C. The solution was replaced with new PBS, and the colon epithelial cells were dissociated by vigorously shaking the tubes. Cells were collected by centrifugation at 1,300 rpm for 5 min at 4 °C. The procedure was repeated 3 times. The cells from all washes were collected, washed once with PBS and used for mRNA isolation.

### Gene expression analysis

Purified colon epithelial cells were homogenized and RNA was isolated by the Trizol method (Ambion, Life Technologies) following the manufacturer’s instructions. Subsequently, the RNA samples were reverse transcribed to cDNA using a Maxima H Minus First Strand cDNA Synthesis Kit (Thermo Fischer Scientific). Relative quantification of gene expression was performed using PowerUp^TM^ SYBR Green Master Mix (Applied Biosystems). QPCR was carried out with initial denaturation at 95 °C for 10 min followed by 40 cycles of denaturation at 95 °C for 15 sec and extension at 60 °C for 1 min. Oligonucleotide primers are shown in Supplementary Table 2. Relative gene expression levels were calculated by ΔΔC_T_ method and normalized to the reference gene β-actin.

### Quantification of butyryl-CoA transferase gene

The quantification of bacterial butyryl-CoA transferase (BCoAT) gene expression levels was measured in bacterial DNA samples from mice under different diets using previously reported primers (Louis and Flint 2007). The data were normalized to 36B4 (36B4F 5’-GCGACCTGGAAGTCCAACTAC – 3’ and 36B4R 5’-ATCTGCTGCATCTGCTTGG – 3’) gene. Higher concentrations of the BCoAT primers (20 µM) were added to each 10 µl reaction in order to account for primer degeneracy (Metzler-Zebeli, Hooda et al. 2010). Each reaction contained 5 ng of faecal DNA and KAPA SYBR master mix (Kapa Biosystems). The amplification program was: 95 °C for 3 min, followed by 40 cycles of 95 °C for 30 seconds, primer annealing at 53 °C for 30 seconds, and 72 °C for 30 seconds of extension.

### Periodic acid-Schiff (PAS) staining

The colons from treated mice were cleaned, prepared as Swiss rolls and frozen in OCT medium. Five-µm sections were prepared in a Cryostat Leica CM1860. The sections were stained with the Periodic acid-Schiff (PAS) staining kit (Sigma), as stated in the protocol. The goblet cells were counted in individual well-preserved colonic crypts.

### Immunofluorescence

Frozen Swiss-rolled colons were cut into 5 µm thick sections and fixed with ice-cold acetone. The tissue was blocked with 3 % BSA in PBST for 1 hour at room temperature and incubated with specific antibodies detecting total β-catenin (H-102), Hsp27 (F-4) and Hsp70 (3A3), all obtained from Santa Cruz Biotechnology. Typically, total β-catenin was used at 1:100 dilution, while Hsp27 and Hsp70 antibodies were used at 1:50 dilution. All primary antibodies were incubated with the tissue at 4°C for overnight. Subsequently, tissue sections were washed 3 times with PBST and incubated with secondary antibodies Goat Anti-Rabbit IgG Alexa-647 (Abcam), and Goat Anti-Mouse IgG Alexa-488 (Santa Cruz Biotechnology) in 1:250 for 1 hour at room temperature. Nuclei were counterstained with DAPI, obtained from Sigma. The tissue was then covered with Fluoromount^TM^ Aqueous Mounting Medium (Sigma) and imaging was performed with a Zeiss LSM510 confocal microscope at the Danish Molecular Biomedical Imaging Center (DaMBIC, University of Southern Denmark).

### Intestinal permeability assay

The intestinal permeability was investigated by oral gavage of the permeability probe FITC-Dextran as described previously (Takahashi, Vereecke et al. 2014). Briefly, 100 µl FITC-dextran (Sigma) dissolved in PBS was administered to mice on a normal diet or 10% LC at a concentration of 600mg/kg body weight. Control mice received 100 µl PBS. After 4 hours, whole blood was collected by cardiac puncture. The serum fraction was isolated by centrifugation at 12,000 rpm for 30 min. Next, three dilutions (1:2.5, 1:5 and 1:10) were made from the serum of each mouse with PBS, and 100 µl of each dilution were added to a 96-well black-walled clear bottom microplate in triplicates together with standards of FITC-Dextran. By using a fluorescence microplate reader (Biotek Synergy H1 multimode plate reader) with an excitation of 488 nm (20 nm bandwidth) and emission wavelength of 528 nm (20 nm bandwidth), the fluorescence of FITC in the serum and the standards was determined. The concentration of the FITC-Dextran in the serum was calculated from the standard curve generated based on the FITC-dextran standards (Supplementary Fig. 1). Finally, the background fluorescence measured in the serum from PBS-treated mice was subtracted from the fluorescence measured in the serum of FITC-Dextran-gavaged animals.

### Determination of endoplasmic reticulum (ER) stress

ER stress was investigated as previously described (Samali, Fitzgerald et al. 2010), with some modifications. Briefly, cDNA from mice under different diets was used as a template for Xbp1 amplification (Xbp1F 5’-GAACCAGGAGTTAAGAACACG–3’ and Xbp1R 5’-AGGCAACAGTGTCAGAGTCC–3’), with β-actin as reference gene. The amplification program was: 95 °C for 3 min, followed by 35 cycles of 95 °C for 1 minute, primer annealing at 62.5 °C for 1 minute, 72 °C for 1 minute and an additional elongation step at 72 °C for 10 minutes. PCR products were separated and analyzed from 2.5% agarose gels. The RT-PCR of all samples was repeated 3 times. In addition, the analysis of the expression of ATF4 and CHOP was carried out by qPCR, as described above in gene expression analysis, and the primer sequences are shown in Supplementary Table 2.

### Statistical analysis

The data were analyzed using GraphPad Prism version 7.0 with unpaired *t*-test tests, as indicated in the figure legends. The statistically significant data correspond to * p ≤0.05, ** p ≤0.01, *** p ≤0.001 and **** p ≤0.0001. Error bars represent SD.

## RESULTS

### A 10% low-carb diet and a normal diet supplemented with 10% apple pectin reduce polyp formation and the proliferation in the colon of Apc^Min/+^Msh2^-/-^ mice

Inactivation of Msh2 in mouse CECs leads to abnormally high proliferation (Belcheva, Irrazabal et al. 2014). Butyrate levels produced from the normal mouse diet were critical for supporting the high proliferation rates and, therefore, sensitised these cells to transformation events. Subjecting Apc^Min/+^Msh2^-/-^ mice to a 7% low carbohydrate diet effectively suppressed polyp formation by reducing butyrate-producing gut microbiota and butyrate levels in the colon of these mice (Belcheva, Irrazabal et al. 2014). These data suggest that altering the luminal concentrations of butyrate is a key factor in controlling cell proliferation and CRC development in this model. Therefore, a better understanding of the interplay between bacterially produced butyrate and cancer-predisposing mutations will be an important step towards designing beneficial dietary regimens.

To gain a further understanding of the effects of differing butyrate concentrations on the proliferation and polyp formation in Apc^Min/+^Msh2^+/-^ (control) and Apc^Min/+^Msh2^-/-^ mice, we used three diets that were expected to generate different amounts of butyrate in the colon. A normal diet (ND) in which 65% of the total calories are derived from carbohydrates is used as a maintenance diet in our mouse facility (Supplementary Table 1). A diet low in carbohydrates, designated as a 10% LC diet, in which metabolized energy from carbohydrates is only 10%, was used to reduce the luminal levels of butyrate. This diet contained 2.3-and 4.3-fold lower amounts of disaccharides and polysaccharides correspondingly and 3.2-fold higher amounts of crude fibre compared to the ND. To increase the production of butyrate above the levels generated by the ND, we supplemented this diet with 10% pectin from apple (10% Pec). Pectin is high in fibre and is known to stimulate butyrate-producing gut microbiota and butyrate production in the colon (Waldecker, Kautenburger et al. 2008, Licht, Hansen et al. 2010). We also treated the mice with the antibiotic metronidazole, which specifically targets butyrate-producing Gram-positive bacteria and was previously reported to reduce polyp numbers in Apc^Min/+^Msh2^-/-^ mice (Belcheva, Irrazabal et al. 2014). All treatments were applied for 3 weeks (Figure 1A), after which the mice were sacrificed. Tissue and faecal content were collected for further analysis. Importantly, the mice on all dietary regimens maintained similar body weight (Supplementary Fig. 2). First, we investigated how these dietary regimens affect the process of polyp formation in Apc^Min/+^Msh2^+/-^ and Apc^Min/+^Msh2^-/-^ mice (Figure 1B). The results showed that the normal diet-fed Apc^Min/+^Msh2^-/-^ mice had a significantly higher number of polyps in the colon compared to their Msh2 WT controls. Metronidazole and a 10% LC diet reduced the polyp numbers. Interestingly, however, ND supplemented with 10% pectin had a similar effect (Figure 1B). Previous work demonstrated that the increased proliferation of CECs from Apc^Min/+^Msh2^-/-^ mice, which is critical for polyp formation, is due to the Msh2 mutation, and APC^Min/+^ has little to no contribution (Belcheva, Irrazabal et al. 2014). Therefore, we investigated whether the above diets reduce polyp development by lowering the proliferation of the CECs in Msh2^+/-^ and Msh2^-/-^ mice. For this purpose, we injected the above-mentioned mice on ND, 10% LC and ND+10% Pec with bromodeoxyuridine (BrdU), a thymidine analogue that is incorporated into the newly synthesised DNA of cells during the S phase. Two hours later, the mice were sacrificed, and the proliferation of the CRCs was analysed from stained colonic sections (Figure 1C). The result showed that the ND-fed Msh2^-/-^ mice had more BrdU-positive cells than their WT Msh2^+/-^ controls (Figure 1C). Both 10% LC and ND+10% Pec diets reduced the percentage of BrdU-positive cells, but only in Msh2^-/-^ mice (Figure 1C). To gain further insight into the effect of these diets on the proliferation of CECs, we assessed the expression of known stem cell markers (Figure 1D). We detected a significant increase in EphB2, EphB3 and CD24 in ND-fed Msh2^-/-^ CECs, in agreement with the previously reported results (Belcheva, Irrazabal et al. 2014). Interestingly, a 10% LC diet significantly reduced all stem cell markers in Msh2^-/-^ CECs. Only CD24 was reduced in Msh2^+/-^ cells. However, the ND+10% Pec regimen significantly reduced only EphB3 mRNA expression levels in Msh2^-/-^ CECs. This result, therefore, indicates that both 10% LC and ND+10%Pec diets decrease the proliferation of CECs and polyp formation in Apc^Min/+^Msh2^-/-^ mice by different mechanisms.

### 10% LC and ND+10% Pec diets induce changes in gut microbiota and production of butyrate in Apc^Min/+^Msh2^-/-^ mice

To further understand the mechanism through which 10% LC and ND+10% Pec diets suppress polyp formation in Apc^Min/+^Msh2^-/-^ mice, we investigated the effects of these treatments on the gut microbiota. It has been previously reported that treatment of mice with metronidazole significantly reduces the total bacterial abundance (Belcheva, Irrazabal et al. 2014). This is also confirmed in this study. However, as a difference from the 7% LC diet that did not affect the total bacterial abundance (Belcheva, Irrazabal et al. 2014), increasing the amount of carbohydrates to 10% in this study significantly decreased this parameter. ND+10% Pec had no effect on the total bacterial abundance (Figure 2A). Furthermore, our results indicate that a 10% LC diet reduced *Clostridium* cluster XIVa, but not *Clostridium* cluster IV (Figure 2B), while supplementation of the ND with 10% Pec did not affect the relative abundance of these bacteria species. Since these two bacterial groups belong to the phylum Firmicutes and are major butyrate-producing species, we further investigated how these dietary treatments impact butyrate production (Figure 2C). Since butyrate diffuses quickly into the CECs and measuring its luminal concentration is challenging, we used an indirect method to quantify butyrate production (Louis, Duncan et al. 2004, Louis and Flint 2007). Butyryl-CoA: acetyl-CoA (ButCoA) transferase has been identified as an enzyme involved in the last step of butyrate synthesis in bacteria (Charrier, Duncan et al. 2006). The use of degenerate primers that amplify the ButCoA transferase gene from genomic bacterial DNA isolated from faecal samples was previously reported as an assay that can estimate the butyrate-producing capacity of the gut microbiota (Louis and Flint 2007). Using this approach, we measured that ButCoA levels in 10% LC were significantly lower compared to the levels measured in ND and ND+10% Pec (Figure 2C). This result correlates with the reduced abundance of *Clostridium* cluster XIVa in 10% LC diet-fed animals (Figure 2B) and suggests that supplementation with 10% Pec did not stimulate butyrate production. Furthermore, one of the best-studied butyrate-responsive genes is the cell cycle inhibitor *p21*. Induction of *p21* expression by butyrate is one of the mechanisms through which butyrate indirectly regulates proliferation in CECs (Archer, Meng et al. 1998). We found that *p21* mRNA levels are significantly lower in Msh2-mutant CECs compared to their WT controls. In addition, the 10% LC diet decreased the mRNA levels of p21 while ND+10% Pec had no effect (Figure 2C). This result, therefore, agrees with the observed changes in the butyrate-producing microbiota and butyrate levels (Figure 2B, C).

**Figure 2.**
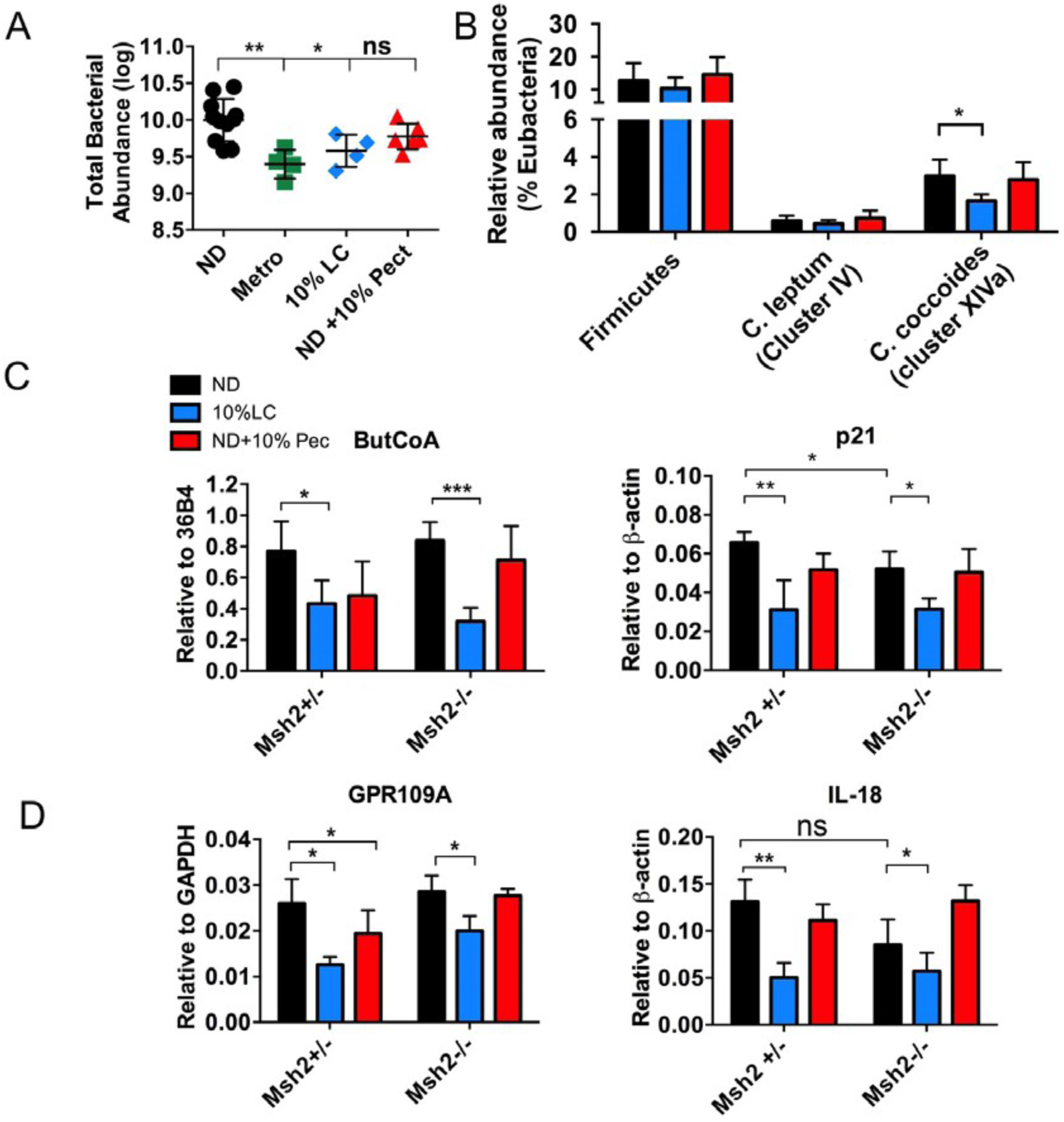
10% low carb (LC) diet and pectin-enriched (ND+10% Pec) diet induce differential changes the abundance of butyrate-producing bacteria and the levels of butyrate in colon. (A) Total bacterial abundance in faeces was calculated by measurement of 16S rRNA gene copies /g sample collected from colons of mice under different treatments. (B) The abundance of Firmicutes and the main butyrate-producing bacterial groups (*Clostridium leptum* and *Clostridium coccoides*), relative to Eubacteria, under different diets was quantified by qPCR analysis. (C) (Left panel) Quantification of butyrate production by determining the expression of the bacterial enzyme butyryl-CoA:acetyl-CoA (ButCoA) transferase by qPCR, relative to 36B4, under the indicated diets. (Right panel) Expression of the butyrate-responsive cell cycle inhibitor p21, relative to β-actin, was assessed by qPCR under different diets in purified colon epithelial cells from Msh2+/-and Msh2-/-mice on the indicated dietary treatments. (D) (Left panel) qPCR analysis for the expression of the butyrate-responsive G-protein coupled receptor GPR109A, relative to GAPDH, in colon epithelial cells from Msh2+/-and Msh2-/-mice under diets. (Right panel) Expression of the GPR109A-produced cytokine IL-18 measured by qPCR, relative to β-actin, in Msh2+/-and Msh2-/-colonic epithelial cells under the indicated diets. Statistical analysis was done by unpaired t-test (n ≥3) and *p*-values are also represented (* *p*-value ≤ 0.05, ** *p*-value ≤ 0.01, *** *p*-value ≤ 0.001).

Butyrate is also shown to activate the G-coupled receptor 109A (GPR109A) and triggers the production of anti-inflammatory cytokine Il-18 that modulates cellular responses to the gut microbiota (Thangaraju, Cresci et al. 2009, Singh, Gurav et al. 2014). We investigated whether GPR109A signalling is influenced by the above-mentioned dietary regimens (Figure 2D). The result showed that GPR109A and Il-18 expression was unchanged in Msh2^-/-^ CECs on ND compared to their WT controls, suggesting that GPR109A signalling does not play a role in the increased proliferation phenotype in these mice. Similar expression of the receptor was also observed in CECs obtained from ND+10% Pec-fed mice. As a difference, a 10% LC diet led to a reduction of GPR109A and IL-18 levels correspondingly, suggesting that reducing the butyrate-producing gut microbiota negatively impacts the GPR109A signalling. Taken together, these results indicate that a 10% LC diet reduces the proliferation of the Msh2-mutant CECs via depleting the levels of butyrate and reduction of the expression of stem cell markers. These data also suggest that ND+10% Pec reduces cell proliferation and polyp formation in Msh2^-/-^ mice by a mechanism that is independent of butyrate.

### 10% LC and ND+10% Pec diets stimulate mucus production in goblet cells

The intestinal epithelium in the colon is covered by a thick layer of mucus, whose main component is Muc-2, produced by goblet cells. The main role of this mucus layer is to protect intestinal cells from direct contact with gut bacteria and their products and it is of great importance for the maintenance of colonic homeostasis (Deplancke and Gaskins 2001, Okumura and Takeda 2017). It has been shown that butyrate induces Muc-2 expression and, thus, protects the CECs against bacteria and derived products in the intestinal lumen (Burger-van Paassen, Vincent et al. 2009). Since a 10% LC diet is associated with decreased production of butyrate, but ND+10% Pec has no such effect, we investigated how these two regimens affect the production of Muc-2 by goblet cells. First, we stained colonic tissue from Msh2^+/-^ and Msh2^-/-^ mice on the indicated dietary regimens with the periodic acid Schiff (PAS) reagent that stains mucins and allows visualisation of the goblet cells (Figure 4A, B). The number of goblet cells per crypt was determined from morphologically well-preserved crypts. The results showed a decrease in the number of goblet cells in Msh2^-/-^ mice under ND compared to their WT controls, in agreement with our previous study (Nøregaard et al., unpublished results). A 10% LC diet reduced the number of goblet cells but only in Msh2^+/-^ mice. Supplementation of the ND with 10% Pec did not affect the goblet cell numbers, regardless of the genotype (Figure 4A, B). Furthermore, we quantified the expression of the goblet cell product Muc-2 and found that it is significantly increased in Msh2^-/-^ mice compared to the Msh2^+/-^ controls. In addition, both dietary regimens dramatically increased Muc-2 expression in both genotypes (Figure 4C). Interestingly, the expression levels of another product of goblet cells, namely Reg-3β, were unaffected by the diets and the genotype (Figure 4C) and suggest that the increase in Muc-2 production observed by both regimens is not dependent on the butyrate levels. Because the changes in the Mu-c2 expression can also be due to specific alterations in the composition of the gut microbiota induced by the respective diets, we further investigated the relative abundance of bacterial groups that are reported to associate with the mucus and either degrade it (Koropatkin, Cameron et al. 2012, Ouwerkerk, de Vos et al. 2013) or stimulate its production (Mattar, Teitelbaum et al. 2002, Caballero-Franco, Keller et al. 2007). We found that both 10% LC and ND+10% Pec significantly decreased the relative abundance of *Lactobacillus* (Figure 3D). 10% LC diet-fed mice had a marked increase in Verrucomicrobia and Bifidobacteria. On the other hand, ND+10% Pec reduced Verrucomicrobia while the abundance of Bifidobacteria was unaffected (Figure 3D). None of the diets affected the relative abundance of *Bacillus*. Interestingly, *Lactobacillus* has been previously reported to positively affect the production of Muc-2 (Mattar, Teitelbaum et al. 2002, Caballero-Franco, Keller et al. 2007). However, our results show that the increased Muc-2 expression is not mediated by *Lactobacillus*, as these species are reduced in both diets. The fact that Muc-2 is significantly increased in ND-fed Msh2^-/-^ colonic tissue compared to their WT controls suggests that the mechanism that upregulates the expression of t h e Muc-2 gene is complex and involves the Msh2 gene inactivation. In addition, the changes in the mucus-associated bacteria that are induced by both dietary regimens (especially observed in Verrucomicrobia and Bifidobacteria) may have an additive effect, further stimulating the expression of Muc-2.

**Figure 3.**
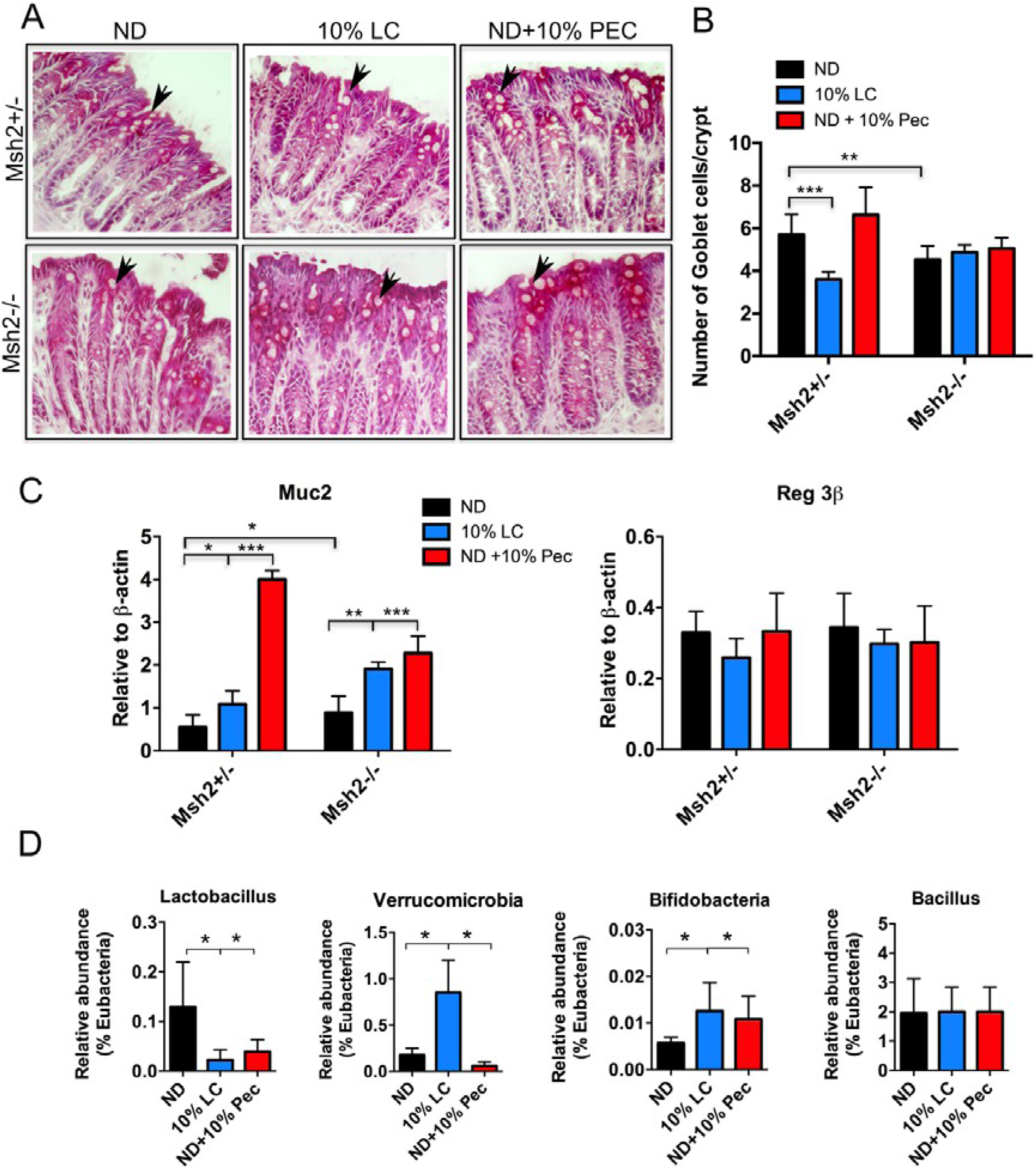
Changes in the Muc-2 expression and mucus associated gut microbiota induced by 10% low carbohydrate diet and normal diet supplemented with 10% pectin. (A) Colonic tissue from Msh2+/-and Msh2-/-mice under different diets was stained with the periodic acid Schiff (PAS) reagent to visualize goblet cells (black arrows). (B) Quantification of the number of goblet cells per colonic crypt was done by calculating the average of goblet cells in, at least, 8 well-preserved colonic crypts in a single mouse (n ≥3 per treatment group and genotype) (C) qPCR analysis of goblet cell-produced compounds mucin-2 (Muc-2) (left panel) and Reg-3β (right panel), relative to β-actin, in Msh2+/-and Msh2-/-mice under different diets. (D) Estimation of the relative abundance of bacterial groups capable of modulating the colonic mucus barrier by qPCR analysis. Data were obtained from at least 6 mice per treatment and analysed by unpaired *t*-test with *p*-values as * *p*-value ≤ 0.05, ** *p*-value ≤ 0.01, *** *p*-value ≤ 0.001.

**Figure 4.**
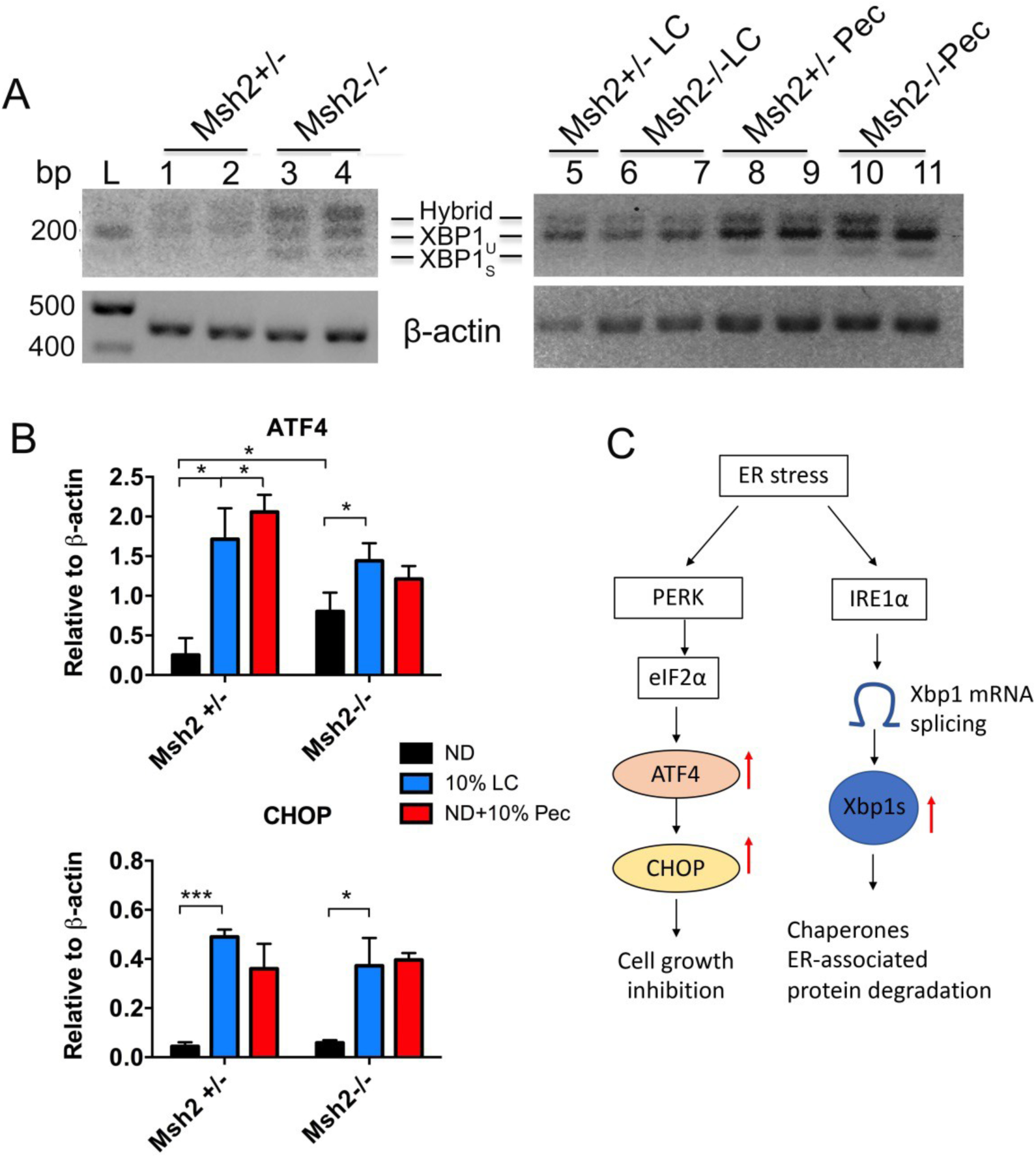
**Induction of Muc2 by 10% LC and ND+10%Pec in Msh2^+/-^ and Msh2^-/-^ mice trigger endoplasmic reticulum (ER) stress**. (A) Analysis of ER stress mediated by IRE-1α pathway. The spliced form of the transcription factor XBP-1 (XBP-1s) was quantified in colon epithelial cells from the indicated mice and dietary regimens by qPCR. The representative images are shown (n ≥3). **(B)** Analysis of ER stress by the PERK pathway. mRNA expression levels of ATF4 and CHOP, relative to β-actin, was carried out by qPCR (n≥4). **(C)** Schematic representation of the IRE-1α and PERK signalling pathways and the mechanisms to resolve cellular ER stress. Red arrows indicate the increased expression of related genes by ND+10% Pec diet. Data were analysed by unpaired *t*-test and *p*-values as * *p*-value ≤ 0.05, ** *p*-value ≤ 0.01, *** *p*-value ≤ 0.001 were also presented.

### Increased Muc-2 production leads to cellular stress in colonic epithelial cells

The observed effects of both 10% LC and ND+10% Pec on the Muc-2 expression are interesting and suggest that these diets that reduce CRC likely affect colonic homeostasis. Muc-2 is a large protein and its proper processing and folding has been associated with stress in the endoplasmic reticulum (ER stress). Therefore, we investigated whether both dietary regimens that increase the Muc-2 mRNA levels cause ER stress. For this purpose, we assessed the activity of two ER stress mediating pathways: IRE1α and PERK. While IRE1α leads to the splicing of the transcription factor XBP1, the PERK pathway activates the transcription of ATF4 and CHOP (Walter and Ron 2011). We used PCR primers that specifically amplify the unspliced (XBP1u) and spliced forms of XBP1 (XBP1s). Both forms of the gene were assessed from agarose gels. The result showed that Msh2^-/-^ CECs had an increase in the spliced XBP1s compared to their WT controls on ND (Figure 4A). This result, therefore, correlates with the increased expression of Muc-2 measured in Msh2^-/-^ cells (Figure 3C). In addition, 10% LC diet-fed mice showed traces of XBP1s, while the spliced form of the gene was abundant in ND+10% Pec-fed CECs, regardless of the genotype. Next, we investigated the expression levels of ATF4 and CHOP in CECs obtained from Msh2^+/-^ and Msh2^-/-^ mice on the respective treatments. First, we found that ATF4 mRNA was increased in Msh2^-/-^ CECs of mice on ND, while CHOP mRNA levels were no different (Figure 4B). Second, both ATF4 and CHOP were elevated by both 10% LC and ND+10% Pec, regardless of the genotype. These results indicate that the increased Muc-2 expression levels observed in ND Msh2^-/-^ as well as under treatment with 10% LC and ND+10%Pec (Figure 3C) correlate with the activation of the ER stress signalling. The results also suggest that under a 10% LC diet, the ER stress is mediated by the PERK pathway, while under ND+10% Pec, both PERK and IRE1α are activated (Figure 4C). Because activation of the ER stress signalling induces the expression of chaperones such as Hsp27 and Hsp70 (Ito, Iwamoto et al. 2005, Mayer and Bukau 2005), we also tested the mRNA expression levels of these genes and found that they dramatically increased by 10% LC and ND+10% Pec, regardless of the genotype (Figure 5A). Furthermore, we stained colonic tissue with specific antibodies for Hsp27 and Hsp70. The results showed that Hsp27 and Hsp70 were overexpressed under the indicated dietary regimens. Both proteins were expressed predominantly at the apical surface (Figure 5B and 5C). The dramatic overexpression of Hsp27 and Hsp70, as well as the increase in Muc-2 expression, further suggest that the intestinal permeability may be altered by these diets. One possible mechanism is the upregulation of the tight junctional protein occludin by Hsp70 (Dokladny, Moseley et al. 2006, Zuhl, Lanphere et al. 2014). To test this possibility, we measured the expression levels of *occludin,* and the result revealed that the gene was dramatically overexpressed under both dietary regimens (Figure 6A). In addition, using FITC-dextran *in vivo* permeability assay, we found that the intestinal permeability of mice on a 10% LC diet was also reduced (Figure 6B).

**Figure 5.**
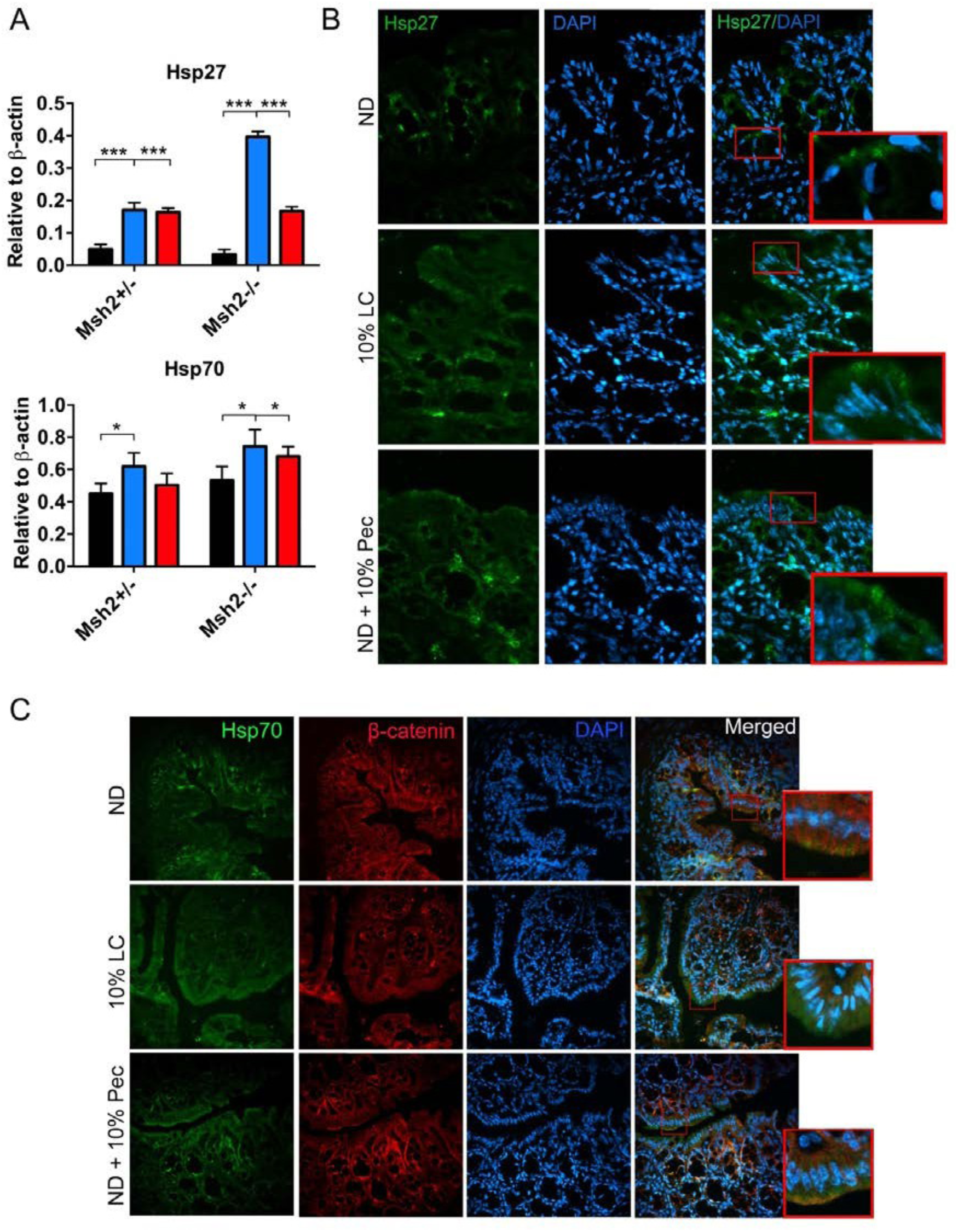
Expression of heat shock proteins Hsp27 and Hsp70 in mice under 10% low carb (LC) and pectin-enriched (ND+10%Pec) diets. (A) qPCR analysis of Hsp27 and Hsp70 in Msh2+/-and Msh2-/-mice under different diets. Data were analysed by (n ≥3) with *p*-values as * *p*-value ≤ 0.05, *** *p*-value ≤ 0.001. (B) Confocal microscopy images of Hsp27 (green) in Msh2^+/-^ mice subjected to different diets. The magnification of the representative images is x40. (C) Confocal microscopy images of Hsp70 (green) and β-catenin (red) in Msh2^+/-^ mice under the mentioned diets. The magnification of the representative images is 20x.

**Figure 6.**
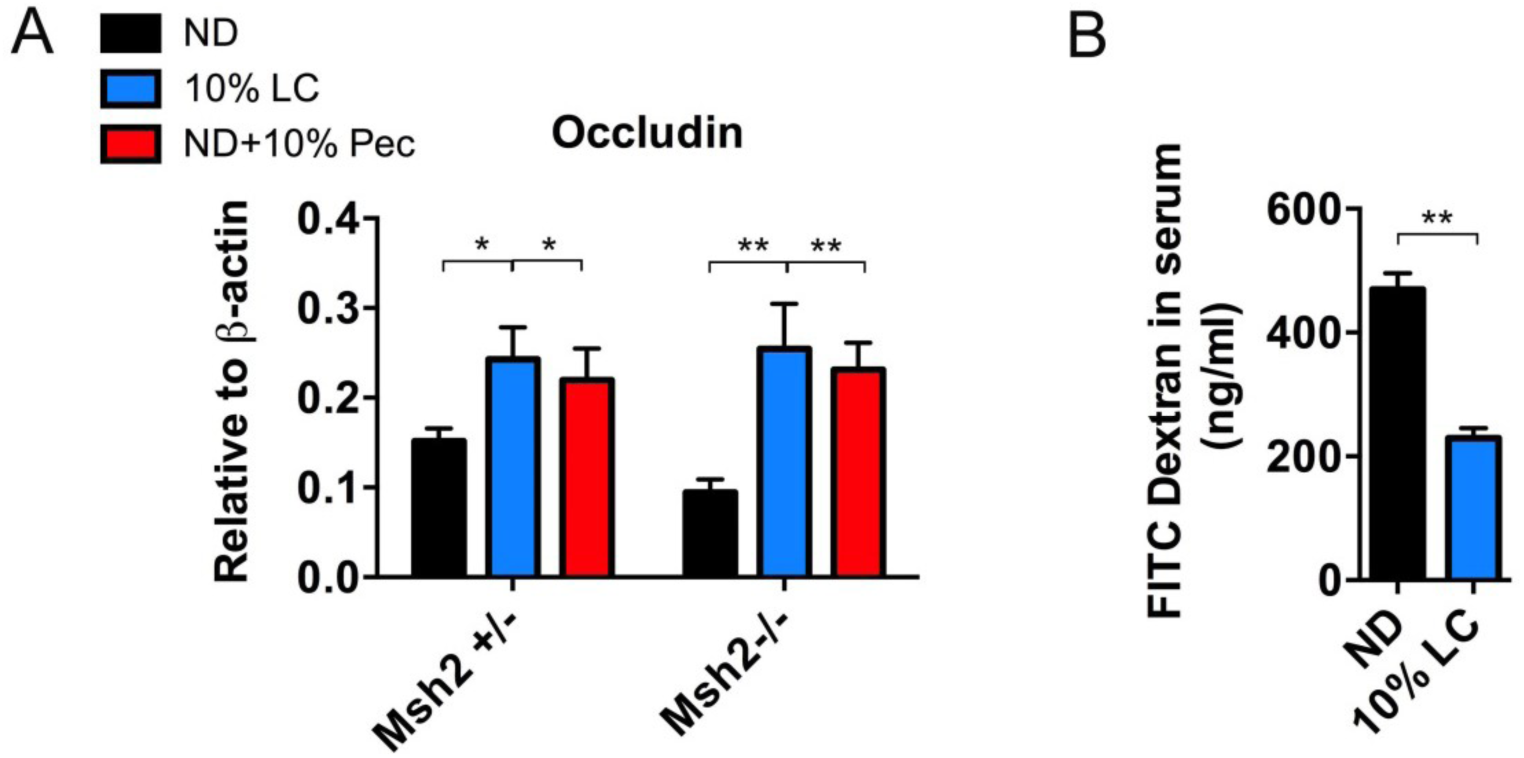
10% LC diet mediates increased barrier function. (A) mRNA expression levels of the tight junctional protein occludin, relative to β-actin, was quantified by qPCR in Msh2^+/-^ and Msh2^-/-^ mice under different diets. (B) *In vivo* permeability assay. The amount of FITC-dextran, in mg/ml, was measured in serum of mice under ND and 10% LC (n ≥4) to determine the intestinal permeability. Data were analysed by unpaired *t*-test and *p*-values (* *p*-value ≤ 0.05 and ** *p*-value ≤ 0.01) were also shown.

Taken together, our results demonstrate that 10% LC and ND+10% Pec diets induce distinct changes in the composition of the gut microbiota and butyrate levels in the colon and that these changes slow the proliferation of the CECs in APC^Min/+^Msh2^-/-^ mice and further reduce polyp formation. Furthermore, the changes in the colonic microenvironment lead to overexpression of Muc-2. As a result, ER stress and expression of Hsp27 and Hsp70 are induced, leading to increased barrier function. These events are likely to protect the cells to cope with the environmental changes. However, the induced cellular stress conditions are known to suppress the proliferation of intestinal cells. Although such an association has to be further investigated, based on these results, it is tempting to propose that a 10% LC diet and ND+10% Pec reduce cell proliferation and polyp development in Apc^Min/+^Msh2^-/-^ mice by triggering cellular stress.

## DISCUSSION

Low carbohydrate (LC) diets have attracted much attention due to their suppressive effect on cancer development. However, the mechanisms through which LC diets can exert this effect are not fully understood. Since CRC initiation and development depend on the complex interactions between genetic mutations, diet, gut microbiota and inflammation defining these mechanisms is essential to novel opportunities for CRC prevention and management.

Previous work demonstrated that 7%LC diet leads to a reduction of bacterially-produced butyrate in the colon and a decrease in the abnormally high proliferation rate and polyp formation in APC^Min/+^Msh2^-/-^ model of CRC (Belcheva, Irrazabal et al. 2014). However, the mechanism through which the 7%LC diet and the low levels of butyrate mediated this effect was not investigated. Indeed, the factors that determine the effects of butyrate either in cancer prevention or in cancer promotion are still not well understood. It has been proposed that the proliferation and metabolic state of the cells (Donohoe, Wali et al. 2012), as well as specific genetic backgrounds (Belcheva, Irrazabal et al. 2014), are central factors that govern the effects of butyrate in CRC. This implies that more studies are needed to further explore the complex interactions between diet, gut microbiota and genetic mutations in CRC. Here we attempted to investigate the effects of two dietary regimens yielding different levels of butyrate on CRC development in the APC^Min/+^Msh2^-/-^ mouse model. We used 10% LC diet that was expected to reduce the butyrate production in the colon by suppressing the growth of the butyrate-producing gut microbiota and normal diet supplemented with 10% Pec (ND+10% Pec) that has been reported to stimulate the growth of the butyrate producers, and hence, to increase the luminal levels of butyrate (Waldecker, Kautenburger et al. 2008, Licht, Hansen et al. 2010). We also treated the mice with metronidazole, which is known to decrease the abundance of butyrate-producing gut microbiota (Belcheva, Irrazabal et al. 2014). This treatment served as a negative control in the study. The results showed that, although not to the same extent, both 10% LC and ND+10% Pec significantly reduced the polyp formation in APC^Min/+^Msh2^-/-^ mice compared to the ND-fed littermates (Figure 1B). In addition, *in vivo* labelling of CECs with BrdU revealed that both regimens reduce cell proliferation (Figure 1C). However, while the 10% LC diet resulted in a decrease in the mRNA levels of the stem cell markers EphB2, EphB3, CD24 and CD44, their expression was little to unaffected by the ND+10% Pec (Figure 1D). This result suggests that both dietary regimens reduced the proliferation and CRC development in APC^Min/+^Msh2^-/-^ mice by different mechanisms. Furthermore, our results demonstrated that the two diets induced different changes in the gut microbiota. The 10% LC diet reduced the total bacterial abundance (Figure 2A) and the relative abundance of *C. coccoides* (cluster XIVa) group (Figure 2B), while ND+10% Pec had no effect. Furthermore, the butyrate production was reduced by 10% LC diet; however, it surprisingly remained unaffected by the supplementation of ND with pectin (Figure 2C). Butyrate is an SCFA that plays a central role in the regulation of colonic homeostasis by being involved in several cellular processes. It is the main energy source for the CECs and regulates their proliferation and immune responses to environmental changes, including changes in the gut microbiota. Furthermore, butyrate has been shown to exert anticancer properties by mediating the cell cycle inhibitor p21 (Archer, Meng et al. 1998). Indeed, we measured a decrease of p21 in Msh2^-/-^ CECs compared to their WT controls. Therefore, lower levels of p21 may be one of the causative factors for the increased proliferation phenotype observed in Msh2^-/-^ cells. We also found that p21 mRNA is decreased in both Msh2^+/-^ and Msh2^-/-^ CECs from mice on 10% LC diet and was not affected by ND+10% Pec (Figure 2C). This result agrees with the measurements of butyrate levels. Since our results show that the proliferation of the CECs is reduced by both regimens, it is not likely that this effect is mediated by p21. Therefore, we explored other mechanisms through which butyrate could affect the proliferation of the CECs.

One mechanism through which butyrate modulates the response of the intestinal epithelium to environmental changes is through the Gpr109A receptor (Singh, Gurav et al. 2014). Indeed, the role of Gpr109A in CRC has just recently emerged. Singh and colleagues have shown that Apc^Min/+^ mice, and the AOM/DSS model of colitis-associated CRC on the Gpr109A^-/-^ background, have increased polyp development (Singh, Gurav et al. 2014). Although the mechanisms through which Gpr109A mediates protection against CRC are not fully understood, it is believed that it integrates key signals from microbial products and coordinates the anti-inflammatory response of the cells. Our results show that GPR109A expression was not different between the genotypes, however, was significantly reduced only in CECs from 10% LC diet-fed mice. This also correlated with low expression of Il-18, suggesting that GPR109A signalling does not play a role in the CRC development of APC^Min/+^Msh2^-/-^ mice.

Another effect of butyrate on colonic homeostasis is its ability to regulate the expression of the Muc-2 gene, and hence, to maintain the barrier integrity. To test whether the altered levels of butyrate in a 10% LC diet are associated with changes in the goblet cell numbers and production of Muc-2, we stained colonic tissue from Msh2^+/-^ and Msh2^-/-^ mice on the ND, 10% LC and ND+10% Pec treatment. Our results show that the number of goblet cells is decreased in ND-fed Msh2^-/-^ mice which is in agreement with our previous work (Nøregaard et al., unpublished results). Interestingly, a 10% LC diet reduced the number of goblet cells but only in Msh2^+/-^ mice (Figure 3A). The number of goblet cells in Msh2^-/-^ mice remained unaffected by the dietary treatments. However, the Muc-2 mRNA was dramatically increased by both the 10% LC diet and ND+10% Pec (Figure 3B).

In addition, the expression of another goblet cell product, namely Reg-3β, was unaffected neither by the diets nor by the genotype (Figure 3C), suggesting that the altered Muc-2 expression is not dependent on the number of goblet cells. A link between gut microbiota, diet and Muc-2 production has been previously described. For example, germ-free mice had smaller and fewer goblet cells in the colon (Comelli, Simmering et al. 2008, Rokhsefat, Lin et al. 2016), and a high-fat diet increased the number of goblet cells together with changes in *Bacteroides*-*Prevotella* abundance (Benoit, Laugerette et al. 2015). Also, Wu and colleagues observed that butyrate and butyrate-producing bacteria increase in Muc-2^-/-^ mice, indicating that the availability of mucus is an important factor that itself shapes the intestinal microbiota (Wu, Wu et al. 2018). One explanation is that the outer layer of the mucus is populated by specific members of the gut microbiota. These species are capable of degrading mucus to use it as an energy source (Koropatkin, Cameron et al. 2012, Ouwerkerk, de Vos et al. 2013). For example, the growth of *Akkermansia muciniphila* that belongs to phylum Verrucomicrobia (Derrien, Vaughan et al. 2004); *Bacteroides fragilis* (Macfarlane and Gibson 1991), and *Bifidobacterium bifidum* (Garrido, Kim et al. 2011) depends on the Muc-2 availability. On the other hand, Lactobacillus species stimulate Muc-2 secretion and, in that way, increase the resistance of the host to pathogens (Mattar, Teitelbaum et al. 2002, Caballero-Franco, Keller et al. 2007). We tested whether the 10% LC diet and ND+10% Pec that affected Muc-2 expression levels will also correlate with changes in the mucus-associated members of the gut microbiota. We found that the relative abundance of Verrucomicrobia and Bifidobacteria was significantly increased in both the 10% LC diet and ND+10% Pec, correlating with increased Muc-2 expression. An increase of Verrucomicrobia was previously reported in fasting hamsters, where the abundance of SCFAs was low (Clarke, Murphy et al. 2012), suggesting that the increase in these species in a 10% LC diet is likely due to the reduced fermentations and production of butyrate. On the other hand, in agreement with our results (Figure 3D), pectin has been shown to stimulate the growth of Bifidobacteria (Ho, Lin et al. 2017, Koutsos, Lima et al. 2017). Interestingly, the mucus-degrading activity of these bacterial species has been shown to stimulate mucus release and increase the thickness of the mucus layer (Caballero-Franco, Keller et al. 2007). Apparently, the regulation of the Muc-2 gene in the colonic epithelium is very complex and depends on different environmental factors, including genetics, the composition of the gut microbiota and availability of butyrate. More studies will need to clarify the molecular details of the Muc-2 expression under different dietary conditions.

However, Muc-2 is a large protein and its proper processing and folding have been associated with ER stress (Tawiah, Cornick et al. 2018). Indeed, our results also confirm this link. Interestingly, we found that under a 10% LC diet, the ER stress is mediated by the activation of the PERK pathway, while under ND+10% Pec both PERK and IRE1α are activated (Figure 4). The ER stress plays an important role in the intestinal homeostasis. PERK signalling is shown to be involved in the normal differentiation of proliferating cells (Heijmans, van Lidth de Jeude et al. 2013). On the other hand, IRE1α activation triggers the splicing of the XBP1 transcription factor that is shown to be important in the maintenance of secretory cells such as goblet cells (Kaser, Lee et al. 2008). ER stress activation ultimately leads to a reduction of cell proliferation (Heijmans, van Lidth de Jeude et al. 2013). This is achieved by inhibition of Cyclin D1 translation, followed by G1 phase cell cycle arrest that provides the cells time to recover from the stress conditions (Brewer, Hendershot et al. 1999). Therefore, the increased levels of Muc-2 likely trigger ER stress that may suppress the proliferation of the CECs.

The activation of ER stress can also induce the expression of chaperones such as Hsp27 and Hsp70 (Ito, Iwamoto et al. 2005, Mayer and Bukau 2005, Heinrich, Tuukkanen et al. 2011). Here we showed that Hsp27 and Hsp70 were dramatically increased by both 10% LC and ND+10% Pec regardless of the genotype (Figure 5). In addition to the ER stress, these proteins can be also induced by many other factors such as butyrate (Ren, Musch et al. 2001, Arvans, Vavricka et al. 2005) or by members of the gut microbiota such as *Bifidobacterium* and *Lactobacillus* (Ueno, Fujiya et al. 2011, Paszti-Gere, Szeker et al. 2012). Importantly, we also measured an increase in Bifidobacterium in both regimens (Figure 3D), suggesting that the changes in the microbiota induced by both regimens may be implicated in the induction of Hsp27 and Hsp70. In addition, L-glutamine, which is increased by 2-fold in the 10% LC diet, is a strong inducer of Hsp70 (Ehrenfried, Chen et al. 1995) and 6% pectin itself has been shown to stimulate Hsp70 (Arvans, Vavricka et al. 2005). The expression of the Hsp27 and Hsp70 may have a different impact on the CECs. They could help the cells to recover from the ER stress (Heinrich, Tuukkanen et al. 2011) or may lead to an increased barrier function via modulating occludin expression (Dokladny, Moseley et al. 2006, Zuhl, Lanphere et al. 2014). Indeed, our results demonstrate that occludin was overexpressed under both dietary regimens (Figure 6A). In addition, this correlated with a marked decrease in the intestinal permeability of mice on 10% LC diet (Figure 6B). Taken together our results demonstrate that a 10% LC diet and ND+10% Pec are associated with distinct changes in the composition of the gut microbiota and butyrate levels in the colon that slow the proliferation of the CECs in APC^Min/+^Msh2^-/-^ mice and reduce polyp formation. However, the changes in the colonic microenvironment induced by these regimens also lead to overexpression of Muc-2. As a result, ER stress and expression of Hsp27 and Hsp70 are induced leading to increased barrier function. These events are likely to protect the cells and help them cope with environmental stress conditions. However, the induced cellular stress has a suppressive effect on the proliferation. Therefore, we speculate that the induced changes in the CECs microenvironment and cellular stress by 10% LC diet and ND+10% Pec ultimately lead to a reduction of cell proliferation and polyp development in Apc^Min/+^Msh2^-/-^ mice. Although further studies will need to prove this link, it raises the question of whether long-term dietary alterations in the colonic microenvironment that protect against CRC are safe for normal colonic homeostasis.

## ACKNOWLEDGEMENTS

We thank Alberto Martin (University of Toronto) for providing the mouse strains used in this study. We also thank Barbara Guerra for her technical help and discussions. The authors would also like to acknowledge the Danish Molecular Biomedical Imaging Center (DaMBIC, University of Southern Denmark) for the use of the bioimaging facilities, as well as Clare Kirkpatrick for the use of the fluorescent plate reader. The project was funded by the Department of Biochemistry and Molecular Biology at SDU.

## AUTHOR CONTRIBUTIONS

M.L.C. designed the study, planned the experiments, conducted data collection and analyses, and wrote the manuscript. A.B. designed and supervised the study, supervised data collection and analyses, and wrote the manuscript.

