## Supplementary information for "Diet-induced changes in the colonic microenvironment reduce polyp development in APC^Min/+^Msh2^-/-^ mice by triggering ER stress"

##### Supplementary figures

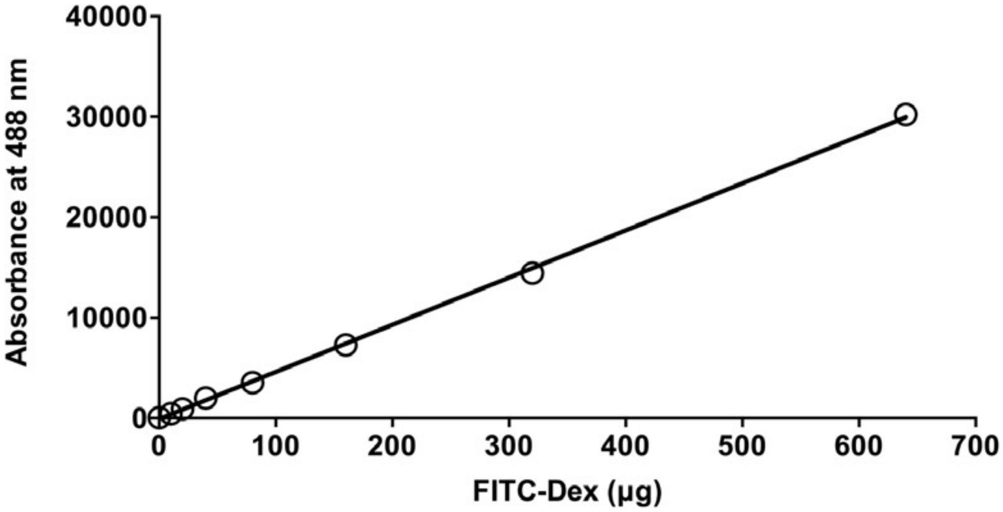


**Supplementary Figure 1. Standard curve of FITC-Dextran (FITC-Dex) for the *in vivo* permeability assay, based on the FITC-Dextrand standards.**

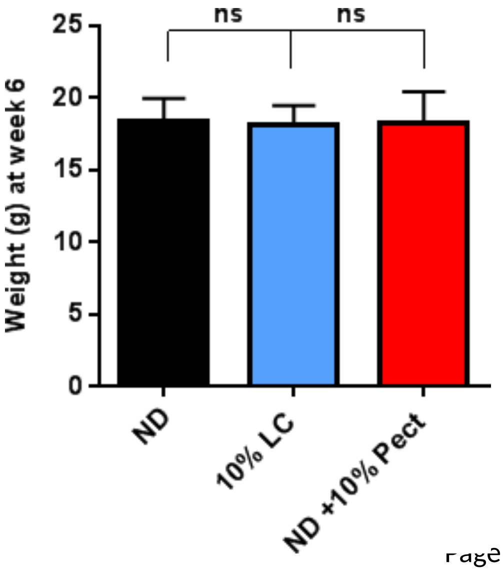

**Supplementary Figure 2. Weight, in grams, of 6-week-old mice under different diets.**

### Supplementary tables

Supplementary Table 1. Overall diet content in the used dietary regimens

| **Ingredients** | **ND** | **10% LC** |
|---|---|---|
| **Protein (%)** | 19.2 | 32.9 |
| **Fat (%)** | 4.1 | 15 |
| **Carbohydrates** | | |
| Disaccharides (%) | 4.8 | 2.1 |
| Polysaccharides (%) | 39.1 | 8.2 |
| Crude fibre (%) | 6.1 | 19.2 |
| Ash (%) | 5.9 | 5 |
| **Energy density, kcal/g** | 3.227 | 3.18 |
| **Calories from protein (%)** | 24 | 75 |
| **Calories from fat (%)** | 11 | 15 |
| **Calories from carbohydrates (%)** | 65 | 10 |

Supplementary Table 2. Quantitative PCR primers for specific bacterial groups

| Bacterial target group | Primer sequence (5'-3') | Annealing Temp (°C) | Refs |
|---|---|---|---|
| *Eubacteria* | ACTCCTACGGGAGGCAGCAGT<br>ATTACCGCGGCTGCTGGC | 60 | (Barman, Unold et al. 2008) |
| *Bacteroides* | GAGAGGAAGGTCCCCCAC<br>CGCTACTTGGCTGGTTCAG | 60 | (Petnicki-Ocwieja, Hrncir et al. 2009) |
| *Firmicutes* | GGAGYATGTGGTTTAATTCGAAGC<br>A AGCTGACGACAACCATGCAC | 54.7 | (Yang, Chen et al. 2015) |
| *Bacillus* | GCGGCGTGCCTAATACATGC<br>CTTCATCACTCACGCGGCGT | 60 | (Petnicki-Ocwieja, Hrncir et al. 2009) |
| *Lactobacillus* | AGCAGTAGGGAATCTTCCA<br>CACCGCTACACATGGAG | 60 | (Rinttila, Kassinen et al. 2004) |
| *C. coccoides*<br>(*C.* cluster XIVa) | ACTCCTACGGGAGGCAGC<br>GCTTCTTAGTCAGGTACCGTCAT | 60 | (Barman, Unold et al. 2008) |
| *C. leptum*<br>(*C.* cluster IV) | CCTTCCGTGCCGGAGTTA<br>GAATTAAACCACATACTCCACTGCTT | 60 | (Furet, Firmesse et al. 2009) |
| *Bifidobacteria* | CGGGTGAGTAATGCGTGACC<br>TGATAGGACGCGACCCCA | 60 | (Furet, Firmesse et al. 2009) |
| *Verrucomicrobia* | TCAKGTCAGTATGGCCCTTAT<br>CAGTTTTYAGGATTTCCTCCGCC | 54.7 | (Yang, Chen et al. 2015) |

Supplementary Table 3. Quantitative PCR primers for stem cell and target genes

| Gene | Primer sequence (5' – 3') | Refs |
|---|---|---|
| EphB2 | GCGGCTACGACGAGAACAT<br>GGCTAAGTCAAAATCAGCCTCA | (Belcheva, Irrazabal et al. 2014) |
| CD44 | CGCTACGCAGGTGTATTCCA<br>TGCTCAGGGCCAACTTCATT | (Belcheva, Irrazabal et al. 2014) |
| CD24 | TTCGCATGGTCACACACTGA<br>ACACACACAGTAGCTTCGGG | This study |
| p21 | GAGCAAAGTGTGCCGTTGTC<br>TCCCAGACAAAGTTGCCCTC | This study |
| p300 | CCCCAAGATCAGCAAATGAAC<br>TGGATAGCCAATTCCTTGGGG | This study |
| GPR109A | TCCAAGTCTCCAAAGGTGGT<br>TGTTTTTCTCTCCAGCACTGAGTT | (Feingold, Moser et al. 2014) |
| Muc-2 | TGTCCTGACCAAGAGCGAAC<br>TTTGAAGGCCACCACGTTCT | This study |
| Reg 3-β | TCCCAGGCTTATGGCTCCTA<br>GCAGGCCAGTTCTGCATCA | (Hogan, Seidu et al. 2006) |
| Hsp27 | AAGGAAGGCGTGGTGGAGAT<br>TTCGTCCTGCCTTTCTTCGT | (Black, Hayden et al. 2011) |
| Hsp70 | CAGCGAGGCTGACAAGAAGAA<br>GGAGATGACCTCCTGGCACT | (Black, Hayden et al. 2011) |
| β-actin | CTACAATGAGCTGCGTGTGG<br>TAGCTCTTCTCCAGGGAGGA | (Xu, Rajesan et al. 2005) |
| ATF4 | ATGGCCGGCTATG-GATGAT<br>CGAAGTCAAACTCTTTCAGATCCATT | (Zhang and Kaufman 2008) |
| CHOP | FCTGCCTTT<br>CACCTTGGAGAC<br>CGTTTCCTGGGGATGAGAT<br>A | (Zhang and Kaufman 2008) |
| IL-18 | CCTCAGATCTTCTGCAACCT<br>TTCCGTATTACTGCGGTTGT | (Singh, Gurav et al. 2014) |
| Occludin | TGGCAAAGTGAATGGCAAGC<br>TTCCTGCTTTCCCCTTCGTG | This study |
